# The ORF8 Protein of SARS-CoV-2 Mediates Immune Evasion through Potently Downregulating MHC-I

**DOI:** 10.1101/2020.05.24.111823

**Authors:** Yiwen Zhang, Junsong Zhang, Yingshi Chen, Baohong Luo, Yaochang Yuan, Feng Huang, Tao Yang, Fei Yu, Jun Liu, Bingfen Liu, Zheng Song, Jingliang Chen, Ting Pan, Xu Zhang, Yuzhuang Li, Rong Li, Wenjing Huang, Fei Xiao, Hui Zhang

**Author notes:** To whom correspondence should be addressed: Hui Zhang. These two authors contribute equally to this work (Yiwen Zhang, Junsong Zhang).

## Abstract

SARS-CoV-2 infection have caused global pandemic and claimed over 5,000,000 tolls^1–4^. Although the genetic sequences of their etiologic viruses are of high homology, the clinical and pathological characteristics of COVID-19 significantly differ from SARS^5,6^. Especially, it seems that SARS-CoV-2 undergoes vast replication *in vivo* without being effectively monitored by anti-viral immunity^7^. Here, we show that the viral protein encoded from open reading frame 8 (ORF8) of SARS-CoV-2, which shares the least homology with SARS-CoV among all the viral proteins, can directly interact with MHC-I molecules and significantly down-regulates their surface expression on various cell types. In contrast, ORF8a and ORF8b of SARS-CoV do not exert this function. In the ORF8-expressing cells, MHC-I molecules are selectively target for lysosomal degradation by an autophagy-dependent mechanism. As a result, CTLs inefficiently eliminate the ORF8-expressing cells. Our results demonstrate that ORF8 protein disrupts antigen presentation and reduces the recognition and the elimination of virus-infected cells by CTLs^8^. Therefore, we suggest that the inhibition of ORF8 function could be a strategy to improve the special immune surveillance and accelerate the eradication of SARS-CoV-2 in vivo.

---

Since the outbreak of COVID-19, the disease has been spreading around the world rapidly^1–4^. Although both COVID-19 and SARS cause severe respiratory illness, the epidemiological and clinical data suggest that the disease spectrum of COVID-19 significantly differ from that of SARS: COVID-19 shows longer incubation period, which is around 6.4 days, ranged from 0 to 24 days; the interpersonal transmissions could occur from pre-symptomatic individuals^5,6^; asymptomatic infection has been widely reported for COVID-19 and severely jeopardize the prevention system in a community^5^; a significant portion of recovered patients still keep shedding viral genetic substances in upper respiratory tract and digestive tract, leading to their stay in the hospital for a significant long time^9–11^; a certain amount of recovered patients turn to re-detectable viral RNA positive after discharge from the hospital^12^. The desynchronization of viral titer and clinical symptom development suggest that the etiologic agent SARS-CoV-2 could have undergone extensive replication in infected host cells without being effectively monitored by host anti-viral immunity.

Cytotoxic T lymphocytes (CTLs) are important for the control of viral infections by directly eradicating the virus-infected cells. In a virus-infected cell, MHC-I molecules present peptides derived from a variety of viral proteins. Once the T cell receptor on CD8^+^ T cells recognizes the special signal presented by MHC-I-peptide complex, the CTL releases various toxic substances including perforins, granzyme, and FasL which directly induce the death of viral-infected cells, as well as many other cytokines such as interferon-γ, TNF-α, and IL-2, etc^8^. As a result, the cells supporting the viral replication will be eradicated and the spread of viruses will be effectively prevented^13^. Some viruses leading to chronic infection, such as human immunodeficiency virus type 1 (HIV-1) and Kaposi’s sarcoma-associated herpesvirus (KSHV), can disrupt antigen presentation for immune envision by down-regulating MHC-I on the surface of cells and evading the immune surveillance^14–16^. Given that SARS-CoV-2 exerts some characteristics of viruses causing chronic infection, we hypothesize that the viral protein(s) of SARS-CoV-2 may affect the antigen presentation system and assist the viruses to escape from immune surveillance.

## Identification of ORF8 as a potent regulator for MHC-I

The genome of SARS-CoV-2 is comprised of ~30,000 nucleotides, sharing 79% sequence identity with SARS-CoV. Similar with SARS-CoV, SARS-CoV-2 has four structure proteins: Spike (S), Envelope (E), Membrane (M), and Nucleocapsid (N)^17,18^. It also harbors some accessory proteins at its 3’ portion (Fig. 1A). Given that the function of almost all structural and non-structural viral proteins of SARS-CoV has been identified, we reasoned that the possible HIV-1 Nef- or Vpu-like function, if exist, would likely fall into the membrane-bound structural proteins or these 3’ accessory ORFs. Initially, we examined the SARS-CoV-2 structural proteins and these un-clarified ORFs for the possible anti-immunity function. Among them, we found that the overexpression of ORF8 in 293T cells indeed significantly down-regulated MHC I (HLA-A2) molecules (Fig. 1B). Specially, the protein sequence of ORF8 of SARS-CoV-2 exhibited the lowest homology with that of SARS-CoV (Fig. 1A)^17–19^. The sequence homology between SARS-CoV-2 and the early-phase SARS-CoV (SARS-CoV_GZ02) in 2003, both of which contain a full-length ORF8, was approximate 26% (Fig. S1A). However, all SARS-CoV strains identified from the mid- and late-phase patients in 2003, such as SARS-CoV_BJ01, contain a 29-nucleotide deletion, resulting in a split ORF8, named as ORF8a and ORF8b respectively (Fig. S1A). The SARS-CoV-2 ORF8 protein was more distant from ORF8a (at 10% sequence identity) and ORF8b (at 16% sequence identity) of SARS-CoV (SARS-CoV_BJ01) (Fig. S1A). Neither the intact ORF8 from SARS-CoV_GZ02, nor ORF8a or ORF8b of SARS-CoV_BJ01 standing for most SARS-CoV strains exerted any effect on downregulating MHC-I (Fig. 1C). The mutation L84S found in the SARS-CoV-2 ORF8 protein was significant for genotyping and phylogenetic analysis^20,21^. However, both L and S subtype of SARS-CoV-2 ORF8 exerted similar effect on down-regulating MHC-I (Fig. 1D).

**Figure 1.**
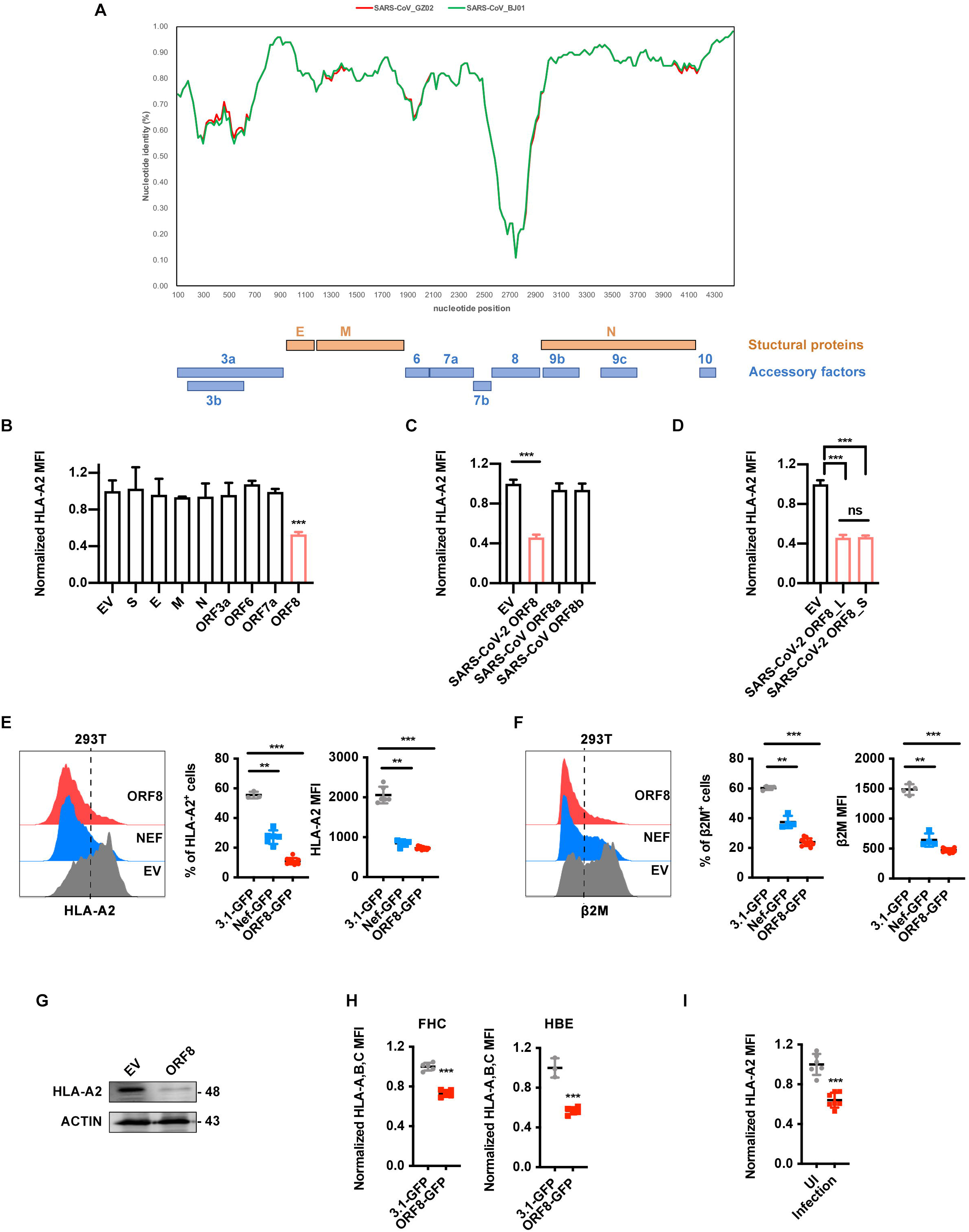
Identification of ORF8 as a potent regulator for MHC-I. **(A)** Similarity plot based on the genome sequence of SARS-CoV-2_WHU01 (accession number MN988668), the genome sequences of SARS-CoV_BJ01 (AY278488) and SARS-CoV_GZ02 (AY390556) were used as reference sequences. The nucleotide position started from the orf3a gene of SARS-CoV-2. **(B-D)** The effect of different viral proteins on the expression of HLA-A2. The viral protein-expressing plasmids were transfected into HEK293T cell line, cells were collected at 48 hours post-transfection for flow cytometry analysis to analyze the MFI of HLA-A2^+^ cells (n=5). The plasmids expressing SARS-CoV-2 structural proteins and ORFs (**B**), SARS-CoV ORF8b and ORF8a **(C)**, and L and S subtype of SARS-CoV-2 ORF8, or empty vector (EV) **(D)** were used. **(E-F)** GFP (negative control), ORF8-GFP, or HIV-Nef-GFP (positive control) expressing plasmid was transfected into HEK293T cells, respectively. Cells were collected at 48 hours post-transfection for flow cytometry analysis. Frequency and MFI of HLA-A2^+^ and β_2_-microglobulin (β_2_M)^+^ cells (gated on GFP^+^ cells) were shown (n=10). **(G)** Western blot analysis for (E) was performed. **(H)** GFP or ORF8-GFP expressing plasmid were transfected into FHC and HBE cells, respectively. Cells were collected at 48 hours post-transfection for flow cytometry analysis (n=5). **(I)** The ACE2 expressing HEK293T cells (HEK293T/Hace2) were infected with SARS-CoV-2 (hCoV-19/CHN/SYSU-IHV/2020) (MOI=0.1). At 48 h post-transfection, cells were collected for flow cytometry analysis (n=5). Data were shown as mean ± SD (error bars). t test and one-way ANOVA was used. P < 0.05 indicates statistically significance difference. * indicates P < 0.05; ** indicates P < 0.01; *** indicates P < 0.001.

In order to further analyze the effect of ORF8 upon downregulating MHC-I, ORF8-expressing plasmid with separate GFP expression (ORF8-IRES-GFP) and control plasmid (3.1-IRES-GFP) were constructed and transfected into 293T cells. A HIV-1-Nef-expressing plasmid (HIV-Nef-IRES-GFP), constructed previously by us, was served as a positive control^22^. The expression of cell surface level of MHC-I heavy chain and the second polypeptide component of MHC-I complex β_2_-microglobulin (β_2_M) was determined by flow cytometry (Fig. S1B). We found that both the frequency and mean fluorescence intensity (MFI) of MHC-I and β_2_M were significantly downregulated by ORF8 overexpression (Fig. 1E and F), as well as the total protein of MHC-I (Fig. 1G). This effect is dose-dependent and increased along with the incubation time (Fig. S1C and D). Furthermore, the MHC-I molecules on various cell lines including human fetal colon cell line FHC, human bronchial epithelial cell line HBE, and human liver cell line Huh7 were also significantly downregulated by ORF8 (Fig. S1E, and Fig1. H). Finally, an authentic SARS-CoV-2 strain named hCoV-19/CHN/SYSU-IHV/2020 were used to infect ACE2-overexpressed 293T cells at MOI 0.1. After 48 h, the infected cells were collected for analysis. We first confirmed the protein expression of ORF8 with Western blot, which is consistent with a recent proteomics data^23^ (Fig. S1F). Importantly, we also found that the expression of MHC-1 on the infected 293T cells was significantly decreased (Fig. 1I).

## MHC-I is targeted for lysosomal degradation by ORF8

To define the mechanism of ORF8-mediated MHC-I reduction, cells were treated with a variety of inhibitors that block membrane protein degradation via different pathways, including N2, N4-dibenzylquinazoline-2,4-diamine (DBeQ), which blocks endoplasmic reticulum-associated protein degradation (ERAD); MG132, which blocks ubiquitin-proteasome system (UPS); and bafilomycin A1 (Baf-A1), which blocks lysosomal degradation. Among these inhibitors, the most significant counteract of MHC-I protein expression reduction by ORF8 was mediated by bafilomycin A1, suggesting that the lysosomal degradation is the major pathway for ORF8-mediated MHC-I downregulation (Fig. 2A and B). Indeed, we found that MHC-I was enriched in lysosomes in ORF8-expressing cells (Fig. 2C). Furthermore, in ORF8-expressing cells, the surface expression of MHC-I was almost abrogated, and redistributed into cytoplasm showing a strong co-localization with LAMP1 (Fig. 2D). We further determined whether ORF8 and MHC-I could interact physically. Through confocal experiment, we first found that ORF8 strikingly co-localized with MHC-I (Fig. 2E). Immunoprecipitation (IP) data further confirmed the binding of ORF8 with either endogenous or exogenous MHC-I (Fig. 2F and G). Collectively, these data suggest that ORF8 can directly bind to MHC-I molecules and targets for lysosomal degradation.

**Figure 2.**
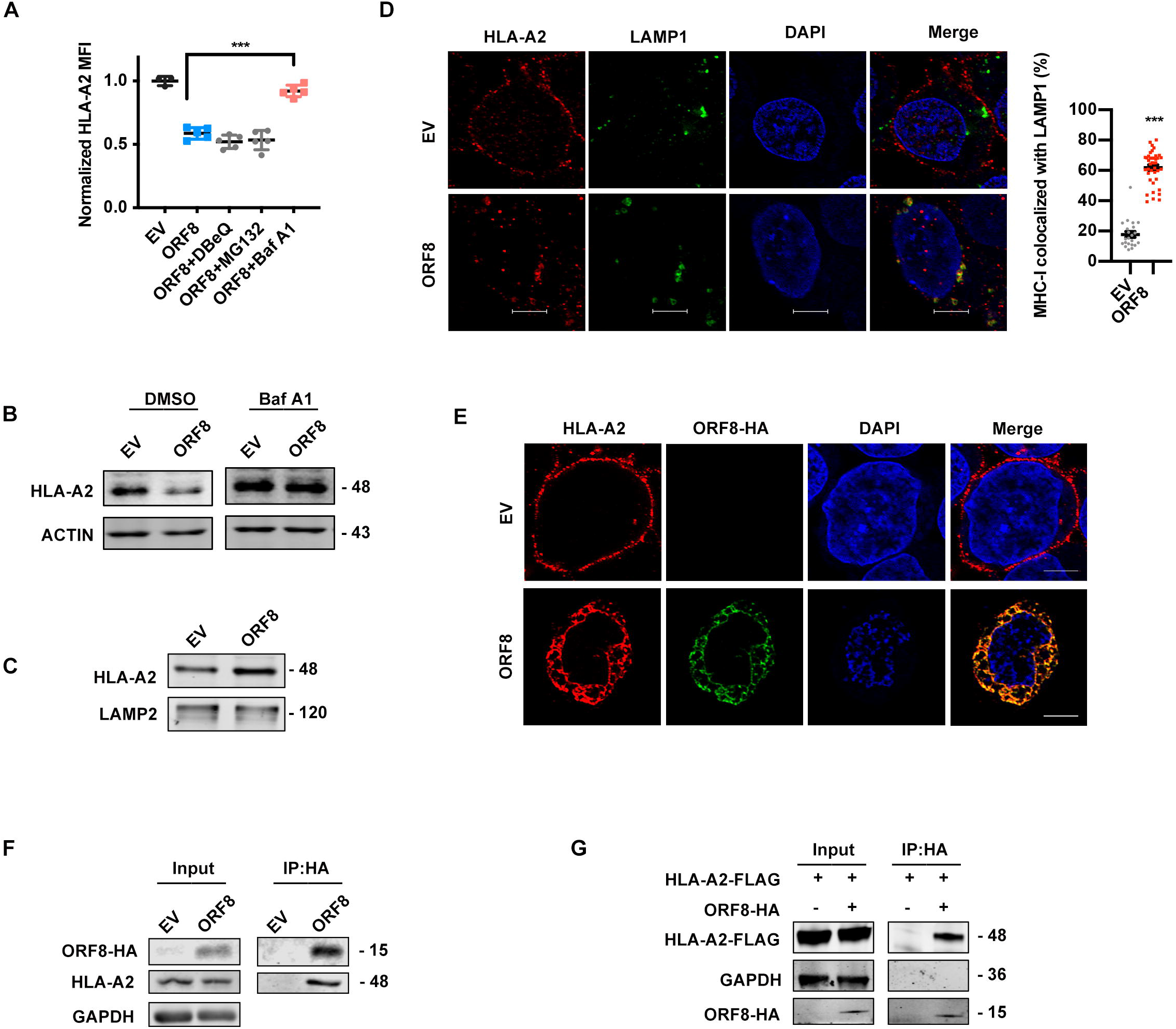
MHC-I is targeted for lysosomal degradation by ORF8. **(A-B)** GFP (EV) or ORF8-GFP expressing plasmid was transfected into HEK293T cells. Before harvest, cells were treated with dimethyl sulfoxide (DMSO), DBeQ (15 μM) for 4 hours, MG132 (10 μM) for 4 hours, bafilomycin A1 (Baf A1, autophagy inhibitor, 100 nM) for 16 hours. The HLA-A2 MFI was analyzed by flow cytometry (gated on GFP^+^ cells) normalized to GFP (EV) group, and the total HLA-A2 protein expression was analyzed by western blotting. **(C)** Cells transfected with empty vector (EV) or ORF8-HA expressing plasmid were treated with Baf A1 (100 nM) for 16 h before harvest for lysosomal fraction. Accumulation of HLA-A2 in lysosomes was analyzed by western blotting. **(D)** Localization of HLA-A2 (red) relative to LAMP1-positive (green) lysosomes. Scale bars, 5μm. Cells were transfected with empty vector (EV) or ORF8-HA expressing plasmid. 24 hours after transfection, colocalization was visualized by confocal microscopy (n=20-40 fields). **(E)** Localization of HLA-A2 (red) relative to SARS-CoV-2 ORF8-HA (green). Scale bars, 5μm. Cells were transfected with empty vector (EV) or ORF8-HA expressing plasmid. 16 hours after transfection, colocalization was visualized by confocal microscopy (n=14-20 fields). **(F)** ORF8 was co-immunoprecipitated with HLA-A2. Empty vector (EV), or ORF8-HA expressing plasmid was transfected into HEK293T cells, respectively. Cells were treated with Baf A1 (100 nM) for 16 h before collected. The cells were treated with cross-linker DSP and co-IP with the anti-HA-tag beads. **(G)** ORF8 was co-immunoprecipitated with the overexpressed HLA-A2. Cells were transfected with HLA-A2-FLAG expressing plasmid together with ORF8-HA expressing plasmid or vector, and treated with Baf A1 (100 nM) for 16 h before harvest. Cells were collected for co-IP with the anti-HA-tag beads. Data were shown as mean ± SD (error bars). t test and one-way ANOVA was used. P < 0.05 indicates statistically significance difference. *** indicates P < 0.001.

## ORF8 mediates MHC-I degradation through an autophagy-dependent pathway

To further identify how SARS-CoV-2 decreases MHC-I expression through ORF8, we performed a mass spectrometry analysis to search for the proteins interacting with ORF8 protein. Consisting with others’ report^24^, the top enrichments of SARS-CoV-2 ORF8 interacting proteins were located at endoplasmic reticulum (ER), indicating that the host interactions of ORF8 may facilitate the significant reconfiguration of ER trafficking during viral infection (Fig. S2A). In addition, ORF8 showed strong co-localization with CALNEXIN^+^ ER and LAMP1^+^ lysosome, rather than GM130^+^ Golgi or RAB5^+^ early endosome (Fig. 3A, Fig. S2B, and Fig. S2C), suggesting that ORF8 most likely performed its function for down-regulating MHC-I at ER or lysosome rather than Golgi or plasma membrane. Also, the knockdown of vesicle-trafficking-related AP1, AP2, or AP3 proteins failed to counteract the MHC-I downregulation mediated by ORF8, further excluding the possible involvements of vesicles transport cargo from the trans-Golgi network, plasma membrane, or endosomal network (Fig. S3A)^25^. Conversely, the knockdown of ERAD-related proteins including HDR1, SEL1L1, ERLIN2, CANX, OS9, or ERLEC1 did not counteract the ORF8-mediated MHC downregulation (Fig. S3B)^26^. The ubiquitination of MHC-I had not significantly changed upon ORF8 over-expression, further excluding the possible involvement of ERAD pathway (Fig. S3C). Thus, we reasoned that ORF8 could mediate MHC-I trafficking from ER to lysosome for degradation.

**Figure 3.**
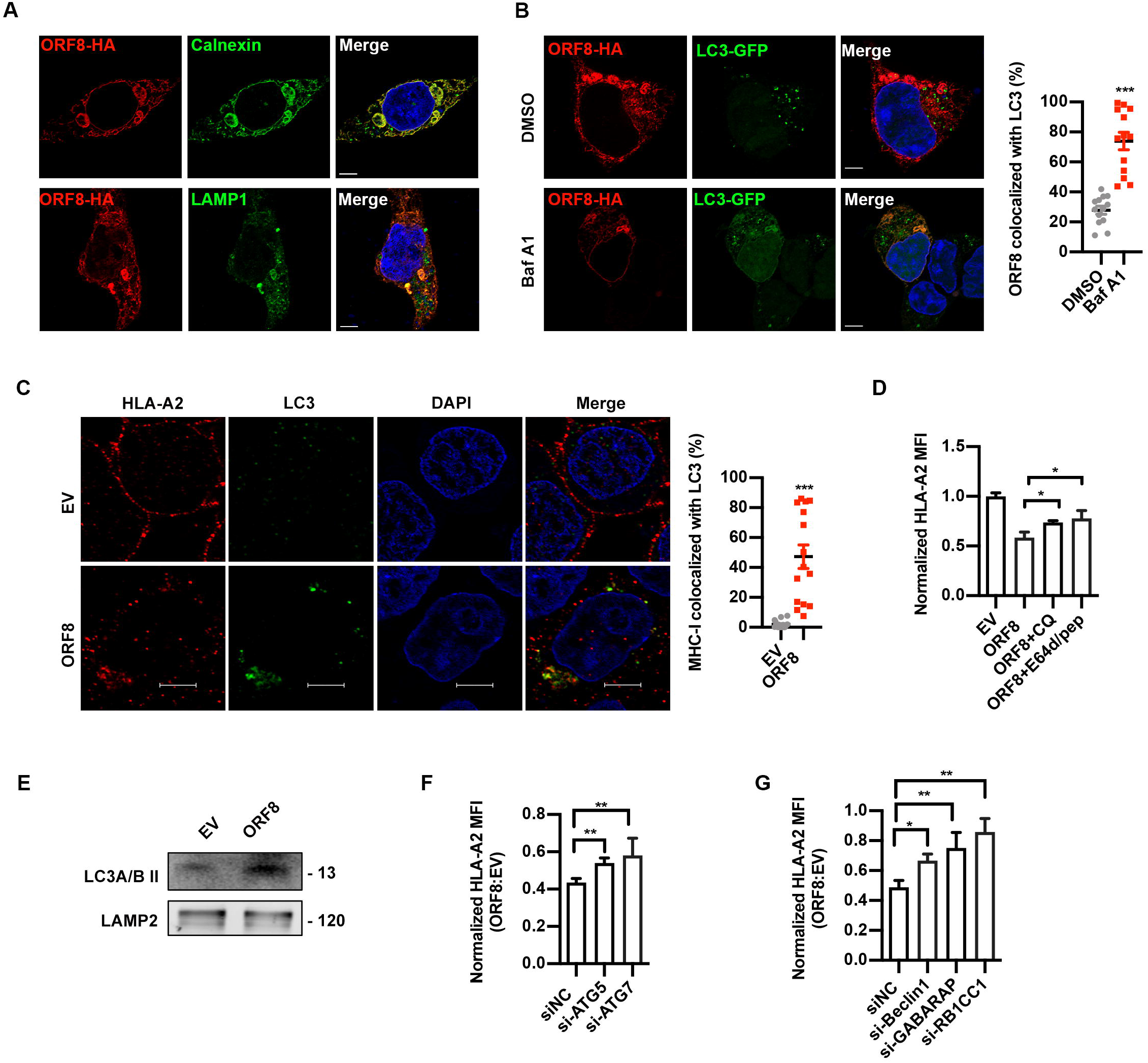
ORF8 mediates MHC-I degradation through autophagy pathway. **(A)** Localization of SARS-CoV-2 ORF8-HA (red) relative to CALNEXIN (green, the top panel) or LAMP1 (green, the bottom panel). ORF8-HA expressing plasmid was transfected into HEK293T cells. 24 hours after transfection, colocalization was visualized by confocal microscopy. Scale bars, 5μm. **(B)** Localization of SARS-CoV-2 ORF8 (red) relative to LC3-GFP (green). ORF8-HA and LC3-GFP expressing plasmids were co-transfected into HEK293T cells. 24 hours after transfection, colocalization was visualized by confocal microscopy. Scale bars, 5μm. (n=14-20 fields). **(C)** Localization of HLA-A2 (red) relative to LC3 (green). ORF8-HA expressing plasmids were transfected into HEK293T cells. 24 hours after transfection, colocalization was visualized by confocal microscopy, Scale bars, 5μm. (n=14-20 fields). **(D)** GFP (EV) or ORF8-GFP expressing plasmid was transfected into HEK293T cells. Before harvest, cells were then treated with chloroquine (CQ) (50 μM) and E64d (10ug/mL) and pepstatin A (pep) (10ug/mL) for 6 hours. The HLA-A2 MFI (gated on GFP^+^ cells) was normalized to GFP group (n=5). **(E)** Empty vector (EV) or ORF8-HA expressing plasmid was transfected into HEK293T cells. Cells were treated with Baf A1 (100 nM) for 16 h before harvest for lysosomal fraction. Accumulation of LC3B in lysosomes was analyzed by western blotting. **(F and G)** GFP (EV) or ORF8-GFP expressing plasmids, and the indicated siRNAs were transfected into HEK293T cells. MFI of HLA-A2 (gated on GFP^+^ cells) was normalized to GFP group (n=5). Data were shown as mean ± SD (error bars). t test and one way ANOVA was used. P < 0.05 indicates statistically significance difference. * indicates P < 0.05; ** indicates P < 0.01; *** indicates P < 0.001.

The trafficking from ER to lysosome is most likely mediated by ER-phagy, which is a kind of selective autophagy and could be divided into three categories: macro-ER-phagy, micro-ER-phagy, and vesicular delivery^27^. Six autophagy cargo receptor proteins bind to and recruit substrates to autophagosomal membranes. To examine their possible involvement, these receptors, including FAM134B, RTN3, ATL3, SEC62, CCPG1, or TEX264, were knocked down with siRNAs respectively^27^. However, we did not observe any effect upon MHC-I expression in the presence of ORF8 (Fig. S3D).

Nevertheless, to search for the possible involvement of autophagy, we first examined the co-localization between ORF8 or MHC-I and autophagosomes within the cells. A substantial fraction of ORF8 co-localized with LC3B-labeled autophagosomes in the ORF8-expressing cells (Fig. 3B). A substantial fraction of the MHC-I puncta also co-localized with LC3B-labelled autophagosomes (Fig. 3C). Furthermore, the specific autophagy inhibitors chloroquine (CQ) and E64/pep restored the expression of MHC-I both on cell surface and total protein level (Fig. 3D, S3E). LC3B was also highly enriched in lysosomes in ORF8-expressing cells (Fig. 3E). Specifically, we found that the knockdown of ATG5, ATG7, Beclin1, and the autophagy cargo proteins RB1CC1 (FIP200) or GABARAP also restored the MHC-I expression, either on the cell surface or cell totality (Fig. 3F-G, S3F-H). However, the knockdown of NBR1 which participates in the MHC-I downregulation in pancreatic cancer cells did not exert any effect, excluding its involvement (Fig. S3I)^28^. Together, although we did yet not observe solid evidence for ER-phagy, at least we found that autophagy plays an important role in ORF8-mediated downregulation of MHC-I.

## ORF8 mediates the resistance of SARS-CoV-2 to antiviral CTLs

It has been known that CTLs participate in immune-mediated protection to coronavirus infection^29^. The consequence of downregulation of MHC-I in the infected cells by ORF8 could be the impairment of CTL-mediated killing of SAR-CoV-2-infected cells. Although no epitope data are yet available SARS-CoV-2, SARS-CoV spike-protein derived peptide-1 (SSp-1, RLNEVAKNL) is predicted to be a potential SARS-CoV-2 epitope^30^, and well characterized for immune response^31^. In order to assess the immune evasion caused by ORF8-mediated MHC downregulation, we generated SSp-1 specific CTLs by sensitization of HLA-A2^+^ healthy donor PBLs with autologous DCs pre-pulsed with SSp-1 (Fig. 4A). SSP-1 pulsed control 293T or ORF8-expressing 293T cells were used as target cells. The result showed the SSp-1 CTLs eliminated ORF8-expressing target cells with lower efficiency (Fig. 4B). Further, we isolated the SARS-CoV-2-specific CD8^+^ T cells from 5 patients who recently recovered from the infection. To evaluate the anti-SARS-CoV-2 activity of CD8^+^ T cells, a mixture of synthetic peptides derived from the S and N proteins of SARS-CoV-2 were used by following the previous SARS studies with minor amino acid modifications^31–33^(supplemental table 1). The PBMCs were treated with the peptide mixture for 7 days, followed by ELISpot analysis to detect IFN-γ secretion. As shown in Figure 4C, compared with healthy donors, the numbers of IFN-γ-secreting CTLs in patient #2, 3 and 5 were much higher than other patients, suggesting that the development of SARS-CoV-2-specific T cell response in these patients. The CD8^+^ T cells of HLA-A2^+^ donor Patient #3 was therefore used for CTL killing assay. The 293T or ORF8-expressing 293T cells were pulsed with synthetic peptide mixture of SARS-CoV-2 and served as target cells, followed by mixture with the effector T cells. Compared with the control, the SARS-CoV-2-specific CTLs also eliminated ORF8-expressing target cells with lower efficiency, indicating that ORF8 protected the target cells from CTL-mediated lysis (Fig. 4D).

**Figure 4.**
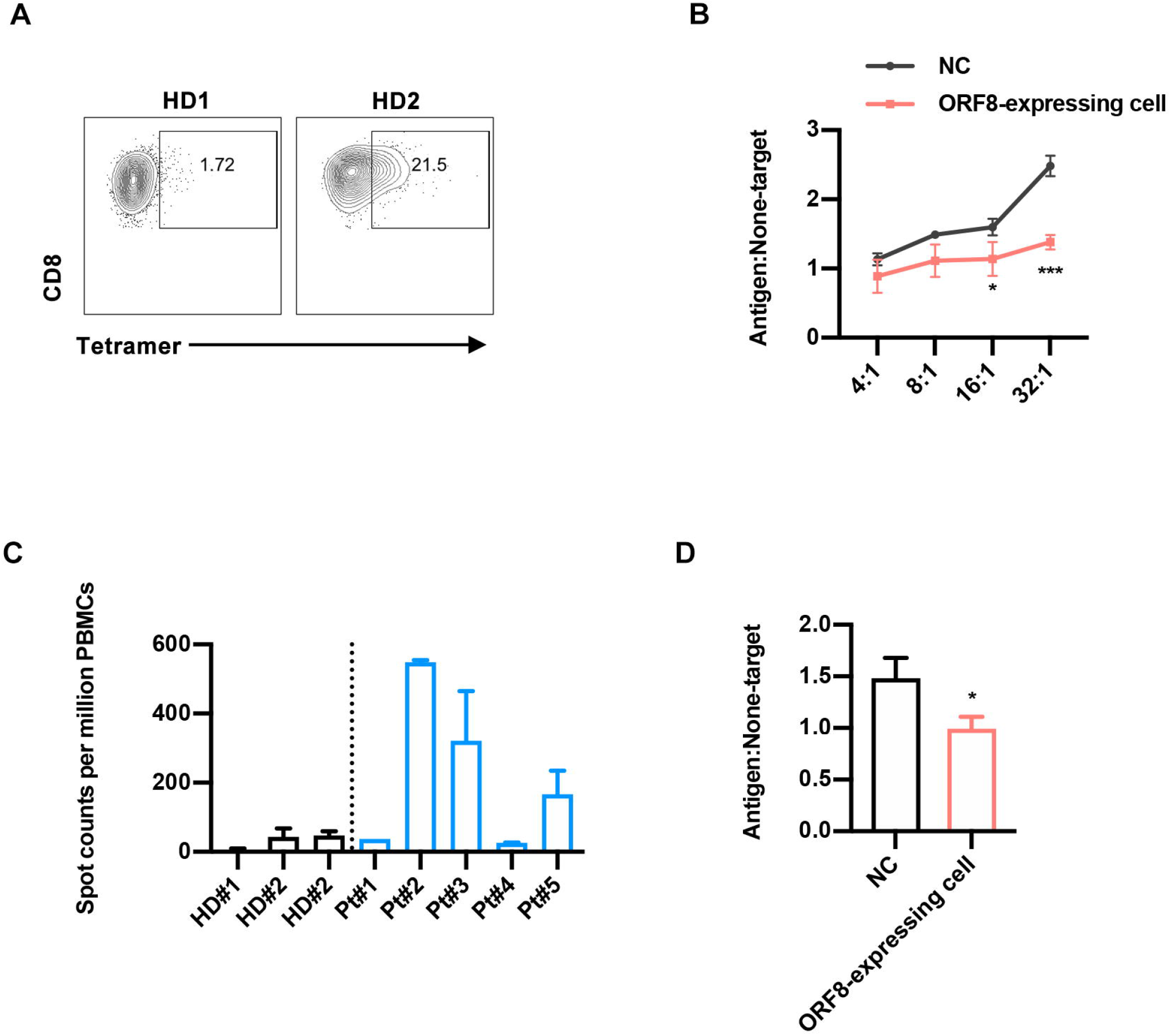
ORF8 mediated resistance of SARS-CoV-2 to antiviral CTLs. **(A)** Frequency of SSp-1-specific CD8^+^ T cells (gated on CD8^+^ cells) generated from HLA-A2^+^ healthy donors (HD) **(B)** Killing assay using SSp-1-specific CD8^+^ T cells generated form healthy donors. CTLs were co-cultured with SSp-1 peptides loaded HEK293T cells (antigen), or with HIV-gag peptide (SL9) loaded HEK293T cells (non-target) overnight. Ratios of dead target versus non-target cells (antigen: non-target) were determined by flow cytometry. **(C)** IFN-γ ELISpot analysis of COVID-19 recover patients (Pt) to synthetic peptides, compared to healthy donors (HD). **(D)** Killing assay using CD8^+^ T cells from HLA-A2^+^ COVID-19 recover patient3. Activated CTLs were co-cultured with SARS-CoV-2 peptides loaded HEK293T cells (antigen), or with HIV-gag peptides loaded HEK293T cells (non-target) at effector: target ratio 8:1. Ratios of dead target versus non-target cells (antigen: non-target) were determined by flow cytometry. Data were shown as mean ± SD (error bars). t test was used. P < 0.05 indicates statistically significance difference. * indicates P < 0.05.

## Discussion

Viral infection would elicit effective innate and adoptive immune response to inhibit the viral replication. Apparently, the anti-viral immunity on SARS-CoV-2 infection remains largely unknown. A proportion of recovered patients still exhibit as the virus carriers and the case of CD8^+^ lymphocytes dysfunction was reported^9–12^. These clinical characters of COVID-19 suggest that SARS-Cov2 could lead to adoptive immune disorder while remain active viral replication. In this report, we have demonstrated that SARS-CoV-2 ORF8 mediates MHC-I downregulation, which is not observed in any other strains of SARS-CoV. The discrepancy of ORF8 between SARS-CoV-2 and SARS-CoV could at least partially be responsible for specific COVID-19 clinical and pathological characteristics, which somehow behaves as a chronic viral infection.

Although some other viruses have also developed the capability to evade the immune surveillance by impairing the antigen presentation, the underlying mechanism is different from each other. HIV-1 Nef mainly facilitates the interaction between AP-1 and MHC and prevent the MHC-I molecule move to plasma membrane. Instead, it re-routes the MHC-I from trans-Golgi network to late endosome/lysosome for degradation^25,34^. K3 and K5 protein of KSHV induce the ubiquitination of MHC-I on the plasma membrane and facilitate its endocytosis^15,35^. Moreover, E3/E19 protein encoded by adenovirus disrupt the association between ER protein TAP and MHC-I and retains MHC-I molecule in ER, impairing the peptide-MHC-I assembly and presentation^36,37^. In this study, we found that ER-resident ORF8 induces the degradation of MHC-I. After excluding the possible involvement of ERAD and other abnormal trafficking, it is reasonably assumed that ER-phagy could participate in this process. However, as we did not find any role played by 6 identified ER-phagy receptors, it is difficult to conclude the involvement of ER-phagy at present^38^. It has been known that ER-phagy is divided into three categories: macro-ER-phagy, micro-ER-phagy, and chaperone-mediated autophagy (CMA). The former is dependent upon the formation of autophagosome^27,39,40^. As we found that the autophagy cargo proteins RB1CC1 (FIP200) or GABARAP are important for ORF8-mediated MHC-I downregulation, it is possible that an unidentified receptor harboring GABARAP-interacting motif (GIM) or FIP200-interacting region (FIR) play a critical role for this process^27,41,42^. Nevertheless, multiple lines of evidence have indicated the participation of autophagy. Based upon our data, we propose that, instead of regular routing through Golgi to plasma membrane, MHC-I at ER is captured by ORF8 and is re-routed to autophagosome and subsequently to autolysosome for degradation (Fig.S4).

SARS-CoV-2 utilize its ORF8 as a unique mechanism to alter the expression of, but not limited to, surface MHC-I expression to evade immune surveillance. Although questions regarding the more detailed molecular mechanisms remain to be further discovered, we provided an important aspect for understanding of how ORF8 disrupts antigen presentation and assistant SARS-CoV-2 immune envision. While current anti-SARS-CoV-2 drugs mainly target enzymes or structural proteins essential to viral replication, our study may promote the development of compounds specifically targeting the impairment of MHC-I antigen presentation by ORF8, and therefore enhancing immune surveillance on SARS-CoV-2 infection.

## Material and Methods

### Ethics statement and patient cohort

This research was approved by the Ethics Review Board of The Fifth Affiliated Hospital of Sun Yat-sen University and Sun Yat-Sen University. The 5 patients who recently recovered from SARS-CoV-2 infection were recruited for this study from The Fifth Affiliated Hospital of Sun Yat-sen University. The given written informed consent with approval of the Ethics Committees were accomplished before the study. Unidentified human peripheral blood mononuclear cells (PBMCs) of healthy blood donors provided by the Guangzhou Blood Center. We did not have any interaction with these human subjects or protected information, and therefore no informed consent was required.

### Cell lines

HEK293T, Huh7 and Vero E6 cell lines was obtained from ATCC. FHC and HBE cell lines are kindly gifted from Professor Wen Liu of Sun Yat-sen University^43^. Theses cell lines conducted authentication through short tandem repeat profiling, karyotyping and cytochrome c oxidase I testing. Test for bacterial and fungal contamination was carried out by using current United States Pharmacopeia methods for viral testing adhering to the United States Code of Federal Regulation (9 CFR 113.53) guidelines, while mycoplasma testing was carried out by direct culture and Hoechst DNA staining and Limulus amoebocyte lysate assay to measure endotoxin values. Cells were maintained in a humidified incubator at 37 °C with 5% CO_2_, grown in Dulbecco’ s modified Eagle’s medium (DMEM) (Gibco) supplemented with 10% FBS (Gibco), 100 units/ml penicillin (Gibco), and 100 μg/ml streptomycin (Gibco).

### Sequence data collection and alignment

The sequences were collected from GenBank database (https://www.ncbi.nlm.nih.gov/nuccore/), including 1 from SARS-CoV-2_WHU01 (accession number MN988668), 1 from SARS-CoV_BJ01 (AY278488), and 1 from SARS-CoV_GZ02 (AY390556). The sequence alignment of complete genome sequences was performed using MAFFT software with default parameters^44^. The protein alignments were created by Clustal Omega software using default parameters conducted in MEGA X^45^. The pairwise sequence identities were calculated using BioEdit software. The similarity analysis based on the genome sequence was performed using SimPlot software^46^.

### Flow cytometry

For analysis of surface markers, cells were stained in phosphate-buffered saline (PBS) containing 0.5% (wt/vol) BSA, with indicated antibodies. Surface proteins were stained for 30 min with the relevant fluorochrome-conjugated monoclonal antibodies and the LIVE/DEAD Fixable Viability Dyes (Thermo Scientific) in PBS containing 0.5% BSA on ice. The following antibodies were used: anti-HLA-A2 (BB7.2), anti-HLA-A,B,C (W6/32), anti-human β2-microglobulin (2M2), and anti-CD8a (53-6.7). Flow cytometry data were acquired on LSR Fortessa (Becton Dickinson).

### Plasmids

The DNA sequences of SARS-CoV-2 structural proteins and ORFs tagged with HA were chemically-synthesized in GENEWIZ (Suzhou, China) and inserted into pcDNA3.1 vector. The ORF8 S mutant expressing plasmid was constructed via a PCR-based mutagenesis method from pcDNA3.1-ORF8-HA by introducing a point mutation (L to S) at the 84 amino acid. The green fluorescent protein (gfp) coding sequence was at the 3’ terminus and constructed into the pcDNA3.1 vector^47^. The IRES-GFP sequence was inserted into 3’ ORF8-HA and named ORF8-GFP. The HIV-Nef-GFP and ubiquitin-HA expressing plasmid was used as previously described by us^22^. All constructs were verified by DNA sequencing. pCMV LC3–GFP was a gift from Dr. Ersheng Kuang of Sun Yat-sen University. The pCMV3-HLA-A-Flag, pCMV-ACE2-Flag, and pCMV3-Rab5-Myc were purchased from Sino Biological. All constructs were verified by DNA sequencing.

### siRNA transfection

siRNAs targeting indicated human genes, and negative control siRNA (siNC) were purchased from RiboBio (Guangzhou, China). Three siRNAs were synthesized for each gene. The siRNAs targeting each gene were transfected as a mixture and have been validated by company to ensure that at least one siRNA was able to knock down target gene mRNA up to 70%. Twelve hours post cell-seeding, cells were transfected with specific siRNAs targeting each genes using Lipofectamine RNAiMAX (ThermoFisher) according to the manufacturer’s instruction. Each gene was set three biological replicates. At 48 h post-transfection, cells were collected for western blot and flow cytometry.

### Infection with authentic SARS-CoV-2

A SARS-CoV-2 strain named as hCoV-19/CHN/SYSU-IHV/2020 strain (Accession ID on GISAID: EPI_ISL_444969) was recently isolated by our lab from a female who was infected at Guangzhou by an Africa-traveler in April 2020 (manuscript submitted to Antimicrobial Agents and Chemotherapy). For infection experiment, HEK293T cells (1.6 × 10^5^ cells/mL) were transfected with pcCMV-ACE2-Flag. After 24 h, cells were washed with PBS and infected with authentic SARS-CoV-2 at MOI=0.1 for 1 h at 37 °C. Then, cells were washed with PBS, and replaced with DMEM (2% FBS). Forty-eight hours after infection, cells were harvested for western blot or testing HLA-A2 expression with flow cytometry.

### Immunofluorescence assay

Immunofluorescence assay was performed as previously described^48^. HEK293T cells were seeded on in μ-slide chambered coverslips (Ibidi; 80826), and transfected as indicated. The transfected cells were treated with DMSO or Baf A1 (100 nM) for 16 h. Forty-eight hours after transfection, cells were washed with PBS and fixed with 4% poly-formaldehyde in room temperature for 10 min, then permeabilized with 0.1% Saponin in PBS for 15 min and blocked with 5% BSA PBS for 30 min. Cells were incubated with primary antibodies at room temperature for 1 h. After washing with 0.1% Tween-20 PBS for three times, cells were stained with secondary antibodies for 1 h, and 4’,6-Diamidino-2-phenylindole dihydrochloride (DAPI) for 5 min. Samples were scanned with Zeiss LSM880 confocal microscopy and analyzed with Imaris. Primary antibodies used in IF assay include anti-GM130 (CST), anti-Calnexin (Proteintech), anti-Rab5 (CST), anti-HA (MBL), anti-Lamp1 (CST), and anti-HLA-A2 (abcam). Images were obtained with LSM880 confocal microscopy (Zeiss). Image analysis and quantification were performed with Imaris 8.4 software (Bitplane).

### Lysosome isolation

For lysosome isolation experiments, HEK293T cells were transfected with indicated plasmids. The cells were treated with 10 μg/mL E64d and 10 μg/mL pepstatin A (pep) for 6 h. Forty-eight hours post transfection, cells were collected for lysosome isolation. The preparation of Lysosomal Fraction was performed by following the manufacturer’s instructions (Sigma, LYSISO1). In brief, the 2.7 PCV of 1× Extraction buffer was added into the cells. The lysis samples were vortexed to achieve an even suspension, and then broken in a 7 mL Dounce homogenizer using Pestle B. Trypan Blue solution staining was used to ascertain the degree of breakage. The samples were centrifuged at 1,000 × g for 10 min and the supernatant was transferred to a new centrifuge tube. The samples were centrifuged again at 20,000 × g for 20 min in microcentrifuge tubes and the supernatant liquid was removed.

### In-cell cross-linking

In-cell cross-linking was performed using dithiobis (succinimidyl propionate) (DSP) (Thermo Scientific) as previously described^49^. DSP were freshly prepared as a 25 mM solution in DMSO and diluted to a working concentration of 0.5 mM in PBS. Cells were washed twice with PBS and then incubated with the cross-linker solution for 30 min at room temperature. Then cells were incubated at room temperature for 15 min with quenching solution (1M Tris-Cl, pH 7.5). Quenching solution was then removed, and cells were washed twice with PBS and cell lysates were prepared for co-immunoprecipitation assay.

### Co-immunoprecipitation (co-IP)

Co-IP assay were performed as our previously described^47^. In brief, HEK293T cells were lysed with NP40 lysis buffer (10 mM Tris-HCl, pH 7.4, 150 mM NaCl, 0.5% NP-40, 1% Triton X-100, 10% glycerol, 2 Mm EDTA, 1 mM Na3VO4, 1% protease inhibitor cocktail (Sigma-Aldrich) and phosphatase inhibitor cocktail (TOPSCIENCE) for 30 min on ice with briefly vertaxing every 10 min. During this period, anti-HA-tag beads were washed three times with ice-cold STN buffer (10 mM Tri-HCl buffered at pH 7.4, 150 mM NaCl, 0.5% NP-40, 0.5% Triton X-100). The lysates were collected and incubated with the prepared anti-HA-tag beads for 4 h or overnight at 4°C with rotating. Then, the immunoprecipitates were washed 4 times with ice-cold STN buffer, eluted by boiling SDS loading buffer, and separated by SDS-PAGE for western blotting or mass spectrometry analysis.

### Mass spectrometry analysis

HEK293T cells were seeded on 10 cm dish and transfected with 12 μg of ORF8-HA, At 48 hours (h) after transfection, cells were collected and lysed for co-IP assay, and the elution was boiled at 100°C with loading buffer supplemented with DTT and separated through 10% SDS-PAGE. The proteins were then visualized with ProteoSilver Plus Silver Stain Kit (Sigma Aldrich) according to the manufacturer’s instructions. The whole lane was cut into ten slices and prepared for liquid chromatography-tandem mass spectrometry (LC-MS/MS) analysis as previously described^48^. Functional pathways representative of each gene signature was analyzed for enrichment in gene categories from the Gene Ontology Biological Processes (GO-BP) database (Gene Ontology Consortium) using DAVID Bioinformatics Resources, observing correlation between two replicate experiments.

### Generation of CTLs in healthy donors

The PBMCs derived from HLA-A2^+^ healthy donors were isolated from peripheral blood by Ficoll-Hypaque gradient separation. PBMCs were resuspended in RPMI 1640 and allowed to adhere to plates at a final concentration of 5×10^6^/ml. After 37°C overnight, non-adherent cells were gently removed. The resulting adherent cells were cultured in medium supplemented with GM-CSF (100 ng/ml, Peprotech) and IL-4 (100 ng/ml, Peprotech) in 5% CO2 at 37°C. Every 2 days, one-half of the medium was replaced by fresh medium containing double concentration of GM-CSF and IL-4 as indicated above. After 5 days of culture, 10 ng/mL recombinant human tumor necrosis factor (TNF-α, Peprotech) was added to the medium to induce phenotypic and functional maturation. Then, 48 hours later, DCs were pulsed with 20μg/mL SSP-1 peptide in the presence of 3μg/mL β-microglobulin (Sino Biological) at 37°C for 3 hours before use. Peripheral blood lymphocytes (PBLs, 2×10^6^) were cocultured with 2×10^5^ peptide-pulsed autologous DCs in a 24-well plate in the presence of 10 ng/mL recombinant human interleukin-2 (IL-2; Peprotech). The next day, recombinant human IL-10 (Peprotech) was added to the culture medium, to give a final concentration of 10 ng/mL. After 7 days, lymphocytes were re-stimulated with peptide-pulsed autologous DCs in medium containing 10 ng/mL IL-2. Lymphocytes were re-stimulated each week in the same manner. At 7 days after the fourth round of re-stimulation, cells were harvested and CD8^+^ cells were purified by microbeads (Miltenyi Biotec) tested by cytotoxicity assay, and tetramer staining.

### Interferon Gamma (IFN-γ) ELISpot

The PBMCs derived from recovered SARS-CoV-2-infected patients were isolated from peripheral blood by Ficoll-Hypaque gradient separation. PBMCs (1×10^6^/ml) were cultured with the synthetic peptide mixture of SARS-CoV-2 (Zhangzhou Sinobioway Peptide Co.,Ltd.). Each peptide was diluted at a final concentration of 1μg/ml in RPMI 1640 medium containing 10% FCS and 20 U/ml recombinant human IL-2 in 24-well culture plate. Half of the medium was changed at day 3 with supplementation of IL-2 at 10 ng/ml. At day 7, IFN-γ-secreting T cells were detected by Human IFN-γ ELISpot assay kits (DKW22-1000-096s; Dakewe) according to the manufacturer’s protocol. PBMCs were plated in duplicate at 4×10^5^ per well and then incubated 24 hours. Spots were then counted using an S6 ultra immunoscan reader (Cellular Technology Ltd.), and the number of IFN-γ positive T cells was calculated by ImmunoSpot 5.1.34 software (Cellular Technology Ltd.). The number of spots was converted into the number of spots per million cells and the mean of duplicate wells plotted.

### Generation of new specificity tetramer using peptide exchange

The peptide exchange experiment was performed with QuickSwitch Quant HLA-A*02:01 Tetramer Kit-PE (MBL) according to the manufacturer’s instructions. Briefly, dissolve each lyophilized peptide (SSp-1) in DMSO at a stock concentration of 10 mM. 50 μl of QuickSwitchTM Tetramer was pipetted into a microtube, and 1ul of target peptide was added and mixed gently with pipetting. Then, 1 μl of peptide exchange factor was added and mixed gently with pipetting. The samples were incubated at least for 4 hours at room temperature protected from light.

### Cytotoxic T lymphocyte (CTL) killing assay

i. For the killing assay for the CTLs generated from healthy donors. CTLs were isolated and counted. A total of 5×10^5^ HEK293T cells transfected with either 3.1-GFP or ORF8-GFP were loaded with 20 μg/ml SSP-1 peptides or HIV-gag peptides (SL9) (at 37 °C for 1 h^31^. The CD8^+^ T cells were cocultured with target cells at the indicated ratios overnight.
ii. For re-stimulation of CD8 T-cells isolated from the recovered SARS-CoV-2-infected patients, the PBMCs were cultured with the synthetic peptide mixture of SARS-CoV-2 at a concentration of 1μg/ml or DMSO in RPMI 1640 medium containing 10% FCS and 20 U/ml recombinant human IL-2 (Peprotech) for 7 days. Then, CTLs from recovered SARS-CoV-2-infected patients were isolated and counted. A total of 5×10^5^ HEK293T cells transfected with either 3.1-GFP or ORF8-GFP were loaded with 20 ug/ml synthetic peptide mixture of SARS-CoV-2 or HIV-gag peptides (SL9) at 37 °C for 1 h^31^. The CD8^+^ T cells were cocultured with target cells. Afterwards, cells were labeled with the fixable viability dye eFluor 780 (FVD, eBioscience) and analyzed by flow cytometry. For determination of antigen: nontarget ratio, cell counts of dead SARS-CoV-2 peptides loaded GFP^+^ cells were divided by the counts for dead HIV-gag peptides loaded GFP^+^ cells^50^.

### Statistical analysis

Differences between two or more groups were analyzed by Student’s t-test or one-way ANOVA followed by Tukey’s test. Statistical significance performed using GraphPad Prism 6. Flow cytometry results were analyzed using FlowJo software (Tree Star Inc.). *P* < 0.05 indicates a statistically significance difference. * indicates *P* < 0.05; ** indicates *P* < 0.01; *** indicates *P* < 0.001.

## Acknowledgments

This work was supported by the National Special Research Program of China for Important Infectious Diseases (2018ZX10302103 and 2017ZX10202102), the Special 2019-nCov Program of Natural Science Foundation of China (NSFC)(82041002), the Important Key Program of NSFC (81730060), and the Joint-Innovation Program in Healthcare for Special Scientific Research Projects of Guangzhou (201803040002) to H.Z.

