## supplemental for "The ORF8 Protein of SARS-CoV-2 Mediates Immune Evasion through Potently Downregulating MHC-I"

### Supplemental Materials

**Supplement Table 1. Synthetic peptides for SARS-CoV-2 used in this work.**

| Position | Sequence | MHC restriction |
|---|---|---|
| S424-433 | KLPDDFTGCV | HLA-A*0201 |
| S1060-1068 | VVFLHVTYV | HLA-A*0201 |
| S976-984 | VLNDILSRL | HLA-A*0201 |
| S996-1004 | LITGRLQSL | HLA-A*0201 |
| S1220-1228 | FIAGLIAIV | HLA-A*0201 |
| S1185-1193 | RLNEVAKNL | HLA-A*0201 |
| S1192-1200 | NLNESLIDL | HLA-A*0201 |
| S449-456 | YNYLYRLF | Unknown |
| N316-324 | GMSRIGMEV | HLA-A*0201 |
| N221-230 | LLLLDRLNQL | HLA-A*0201 |
| N345-353 | NFKDQVILL | Unknown |
| M89-97 | GLMWLSYFI | HLA-A*0201 |
| M148-156 | HLRIAGHHL | HLA Class I |

Fig S1

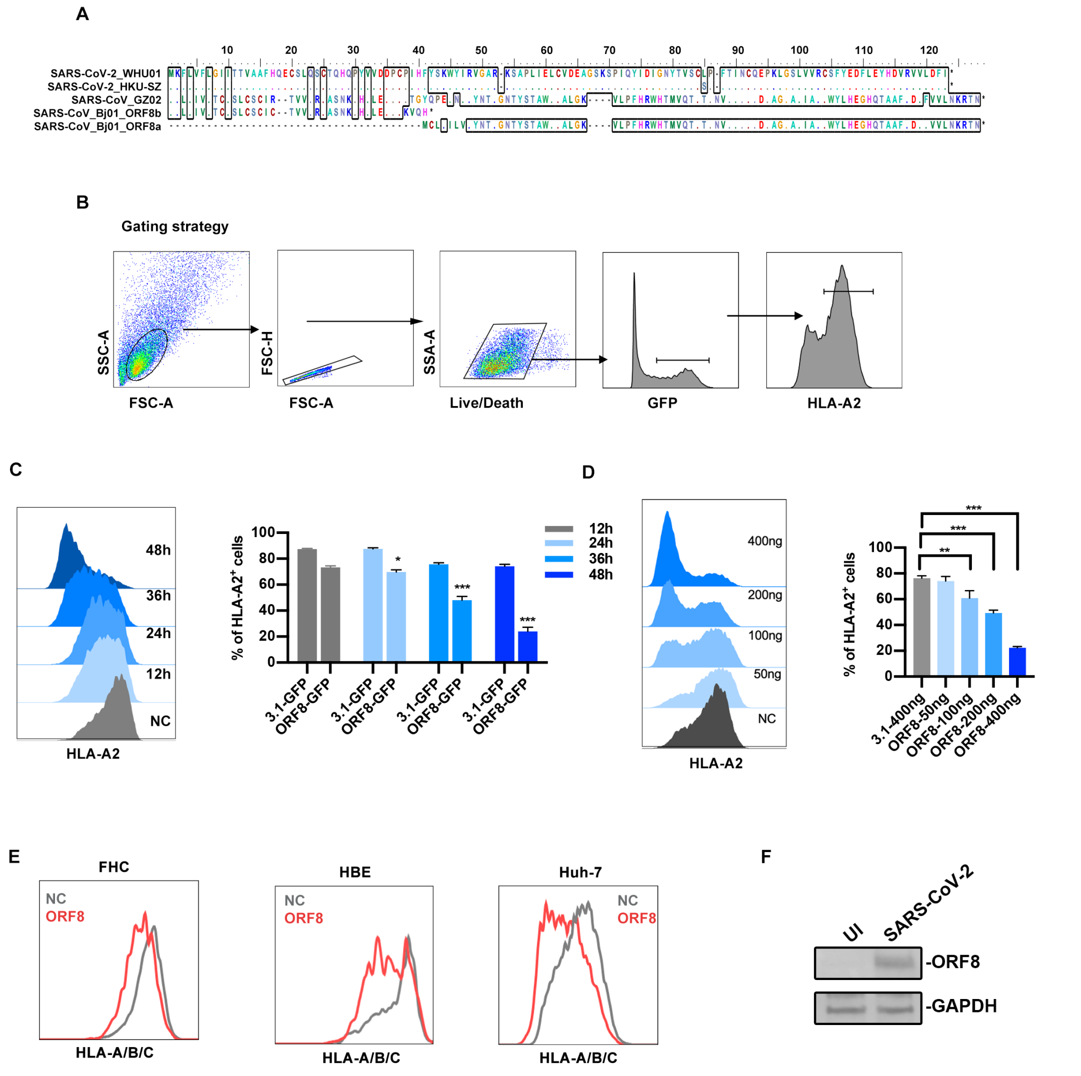


**Supplement Fig. 1 Surface expression of MHC-I is downregulated in ORF8-expressing cells. (A)** The multiple alignment was created based on the amino acid sequences of the encoded protein of orf8 gene, including 1 from SARS-CoV-2_WHU01, 1 from SARS-CoV-2_HKU-SZ with L84S mutation, 1 from SARS-CoV_GZ02, and 1 from SARS-CoV_BJ01 orf8a, 1 from SARS-CoV_BJ01 orf8b. The similarity shading with the color was referred to the chemistry of each amino acid at that position. **(B)**

Gating strategy for FACS analysis of MHC-I expression used in this study. **(C)** GFP, or ORF8-GFP expressing plasmid was transfected into HEK293T cells, respectively. Cells were harvested for flow cytometry analysis at indicated time point. Frequency of HLA-$A2^+$ cells (gated on $GFP^+$ cells) were shown (n=5). **(D)** Different doses of GFP, or ORF8-GFP expressing plasmid were transfected into HEK293T cells, respectively. After 48 hours cells were harvested for flow cytometry. Frequency of HLA-$A2^+$ cells (gated on $GFP^+$ cells) were shown (n=5). **(E)** GFP, or ORF8-GFP expressing plasmid was transfected into FHC, HBE or Huh7 cells, respectively. At 48 hours post-transfection, cells were harvested for flow cytometry analysis (n=5). **(F)** The ACE2 expressing HEK293T cells (HEK293T/Hace2) were infected with SARS-CoV-2 (hCoV-19/CHN/SYSU-IHV/2020) (MOI=0.1). At 48 h post-transfection, cells were collected and followed by western blotting. Data were shown as mean ± SD (error bars). t test and one way ANOVA was used. $P < 0.05$ indicates statistically significance difference. * indicates $P < 0.05$.

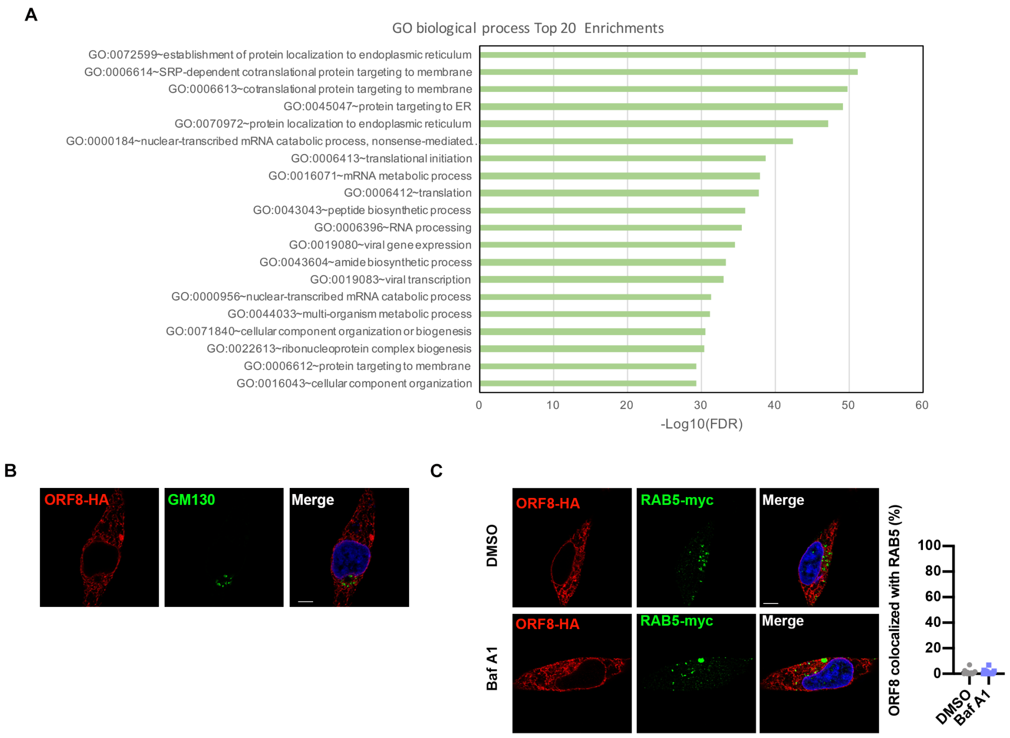


**Supplement Fig.2 ORF8 interacting with endoplasmic reticulum (ER). (A)** The top 20 significant enrichments of the gene ontology (GO) biological process terms. Transformed false discovery rate (FDR) was indicated at the X-axis. **(B)** Localization of SARS-CoV-2 ORF8 (red) relative to GM130 (green, the top panel). ORF8-HA expressing plasmid was transfected into HEK293T cells. Cells were treated with DMSO or Baf A1 (100 nM) for 16 h. At 24 hours after transfection, co-localization was visualized by confocal microscopy. Scale bars, 5μm. (C) Localization of SARS-CoV-2 ORF8 (red) relative to RAB5 (green). ORF8-HA expressing plasmid together with RAB5-myc expressing plasmid expressing plasmid were transfected into HEK293T cells. Cells were treated with DMSO or Baf A1 (100 nM) for 16 h. At 24 hours after transfection, co-localization was visualized by confocal microscopy. Scale bars, 5μm, (n=14-20 fields).

Fig. S3

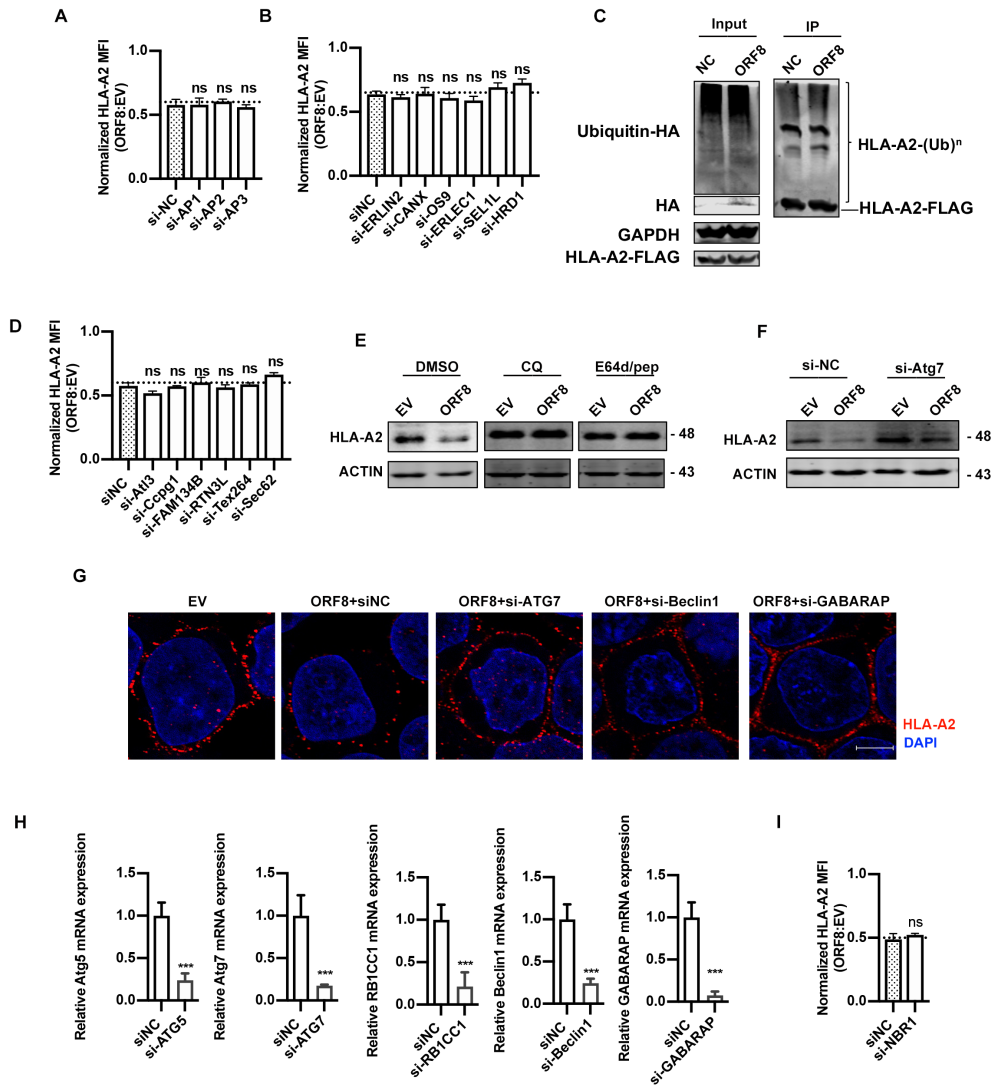


**Supplement Fig.3 ORF8 mediates MHC-I trafficking from ER to lysosome for degradation. (A-B)** GFP (EV) or ORF8-GFP expressing plasmid, and indicated siRNAs was transfected into HEK293T cells. MFI of HLA-A2 (gated on $GFP^+$ cells) was normalized to GFP group (n=5). **(C)** HLA-A2-FLAG and ubiquitin-HA expressing plasmids together with ORF8-HA expressing plasmid or empty vector were transfected into HEK293T cells. Cells were treated with MG132 (10μM) for 12 h before harvest. HLA-A2 ubiquitination was analyzed by Co-IP with anti-Flag-tag beads and followed by western blotting. **(D)** GFP (EV) or ORF8-GFP expressing plasmid, and indicated siRNAs were transfected into HEK293T cells. MFI of HLA-A2 (gated on $GFP^+$ cells) was normalized to 3.1-GFP (n=5). Data were shown as mean ± SD (error bars). t test was used. **(E)** GFP (EV) or ORF8-GFP expressing plasmid was transfected into HEK293T cells. Before harvest, cells were then treated with chloroquine (CQ) (50 μM) and E64d (10ug/mL) and pepstatin A (pep) (10ug/mL) for 6 hours. The total HLA-A2 protein expression was analyzed by western blotting. **(F)** GFP (EV) or ORF8-GFP expressing plasmids, and the indicated siRNAs were transfected into HEK293T cells. The total HLA-A2 protein expression was analyzed by western blotting. **(G)** ORF8-HA expressing plasmids, and the indicated siRNAs were transfected into HEK293T cells. At 48 hours after transfection, HLA-A2 localization was visualized by confocal microscopy. (n=14-20 fields). **(H)** Indicated siRNAs were transfected into HEK293T cells. At 48 hours after transfection, the siRNA efficiency was measure by realtime qPCR (n=5). (I) GFP (EV) or ORF8-GFP expressing plasmid, and indicated siRNAs was transfected into HEK293T cells. MFI of HLA-A2 (gated on $GFP^+$ cells) was normalized to GFP group (n=5). P < 0.05 indicates statistically significance difference. * indicates P < 0.05.

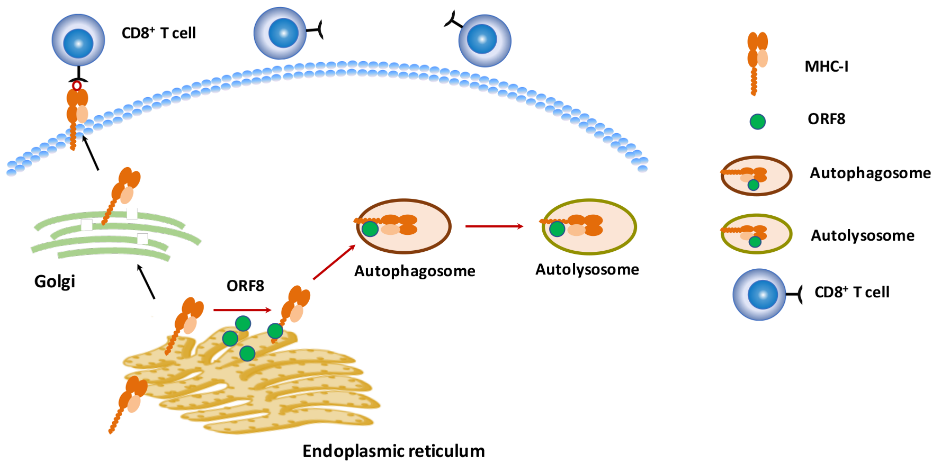


**Supplement Fig.4 Schematics for that ORF8 mediates MHC-I lysosome degradation through an autophagy-dependent pathway.**
